# Functional traits and phylogeny predict vertical foraging in terrestrial mammals and birds

**DOI:** 10.1101/2024.04.18.589860

**Authors:** Patrick Jantz, Andrew Abraham, Brett Scheffers, Camille Gaillard, Mike Harfoot, Scott Goetz, Christopher E. Doughty

## Abstract

Earth’s ecosystems are characterized by numerous gradients related to the distribution of environmental conditions and resources. Niche theory predicts that animals will evolve traits to exploit changing resource availability and environmental conditions across these gradients. Much work has been done examining how animal traits like body mass and diet change across gradients from regional to global scales. Environmental and resource gradients in the vertical dimension tend to exhibit strong changes over relatively short distances due to the influence of elevation and vegetation. Vegetation structure may be an especially important vertical axis as it contributes to strong gradients in micro- climate, food resources, and predation risk. To investigate interrelationships between the vertical niche and its presumed drivers, we use functional traits, phylogenies, and predation risk to predict the vertical foraging niche for 4,828 mammals and 9,437 birds globally. To provide biogeographic context to the predictive analysis, we use species ranges to map geographic distributions of the vertical foraging niche and relationships between the niche and its presumed drivers. Linking trait databases with species range maps revealed distinct global distributions of vertical foraging niches for mammals and birds. The most important predictors of these niches varied by taxon but there were several systematic relationships. Diet, body mass, and phylogeny were strong predictors of vertical foraging niche across mammal and bird species. Percent fruit in diet exhibited progressively more positive relationships with higher canopy foraging positions. Predation pressure was relatively unimportant in predicting most vertical foraging niches for birds and mammals but displayed a positive trend with arboreal foraging. Geographic hotspots for the importance of fruit in both mammal and bird diets included the Andes-Amazon transition zone, the Amazon Basin, and New Guinea. Our results provide support for the theory of resource driven vertical niche partitioning but also reveal that vertical niches are strongly associated with phylogeny, suggesting niche conservatism in numerous mammal and bird families. Geographic patterns in variable importance values suggest multiple mechanisms behind spatial structure in eco-evolutionary relationships, including latitudinal gradients in vegetation structure and composition, historical patterns of island isolation (in Southeast Asia), and the influence of habitat heterogeneity driven by tectonic processes (in South America).

## Introduction

Earth’s ecosystems are characterized by numerous gradients related to the distribution of environmental conditions and resources. Niche theory predicts that animals will evolve traits to exploit changing resource availability and environmental conditions across these gradients (Gámez and Harris, 2022; McGill et al., 2006). Much work has been done examining how animal traits like body mass and diet change across gradients from regional to global scales. One of the most well recognized patterns is Bergmann’s rule, where animal body mass increases towards the poles (Blackburn et al., 1999).

Environmental and resource gradients in the vertical dimension tend to exhibit strong changes over relatively short distances due to the influence of elevation and vegetation (Doughty et al., 2017; Janzen, 1967; Nakamura et al., 2017; Xing et al., 2023). For example, each dietary guild of passerine birds supports many more species in lowland portions of an Andes-Amazon elevation gradient than it does in high elevation portions, corresponding to large changes bird species richness over a short distance (Pigot, 2016).

Vegetation structure may be an especially important vertical axis as it contributes to strong gradients in micro-climate (e.g., temperature, (Nakamura et al., 2017)), resources (e.g., high energy fruits, (Fleming et al., 1987; Schaefer et al., 2002), and risk (e.g., predation, (Shattuck and Williams, 2010).

Consequently, some of the wide array of functional traits possessed by animals likely evolved to exploit ecosystem resources across vegetation vertical gradients, thereby determining the types of ecosystem resources that are available to a given animal as well as the influence that animals may have on ecosystem structure and function (Lundgren et al., 2021; Oliveira and Scheffers, 2019).

In addition to functional traits, shared evolutionary history, indicated by phylogenetic relationships, may also explain patterns of vertical niche use. The emergence of vegetation types that contribute to contemporary patterns of vegetation structure occurred several million years ago (Tiffney, 1984), providing opportunities for the evolution and radiation of groups to exploit this structure (Fleming et al., 1987) as well as opportunities for niche shifts (Bolmgren and Eriksson, 2005). Where phylogeny is an important predictor of vertical niche use, it suggests that related species possess a set of conserved traits, body shape and limb length, for example (Linden et al., 2023) that allowed those species to occupy a particular vertical niche over long time scales (Wiens et al., 2010).

Trophic interactions may also vary with vegetation structure. Theories of animal behavior suggest that predation interacts with vegetation structure to contribute to a “landscape of fear”, where predation risk varies according to landscape context and predator distribution, influencing habitat use by prey species (Laundre et al. 2010). Studies on arboreal and scansorial (climbing) species show that they respond to the landscape of fear in three dimensions, reducing threats from ground hunting predators via escape into the canopy (Emerson et al., 2011; Makin et al., 2012; McGraw and Bshary, 2002).

To investigate interrelationships between the vertical niche and its presumed drivers, we use functional traits, phylogenies, and predation risk to predict the vertical foraging niche for 4,828 mammals and 9,437 birds globally. To provide biogeographic context to the predictive analysis, we use species ranges to map geographic distributions of the vertical foraging niche and relationships between the niche and its presumed drivers.

## Methods

### Functional Traits

Traits that represent an organism’s functional niche in the ecosystem such as diet, body mass, and activity period, so called “Elton Traits” (Wilman et al., 2014), represent a promising candidate set for explaining vertical niche use. We obtained information on diet, foraging stratum, body mass, and activity period for mammal and bird species from the EltonTraits 1.0 database (Wilman et al., 2014). In EltonTraits, mammals are classified by foraging stratum into marine, ground (including aquatic foraging), scansorial, arboreal or aerial types. For birds, the database provides estimates of proportional use of several foraging strata: below the surface of the water, at the surface of the water, on the ground, in the understory, at mid to high levels in trees or in high bushes but below the top of the canopy, in or just above the top of the canopy, or aerially. Vertical foraging niches for mammals and birds serve as the response variable in the models described below. In the construction of EltonTraits, traits for species with little or no information were imputed from genus level, or rarely, family level typical values (Wilman et al., 2014). We conducted sensitivity tests when predicting vertical foraging niche by removing species with imputed values and comparing with predictions made for species with imputed values. For both birds and mammals, we filtered the database to exclude marine and pelagic species.

We used the Global Biodiversity Information Facility’s taxonomic backbone (GBIF Secretariat 2021) to standardize species names and facilitate matching with other databases. For modeling purposes, we represented mammal foraging in each stratum as a binary variable and maintained representation of bird foraging in each stratum as a percent variable (Table 1).

**Table 1.**
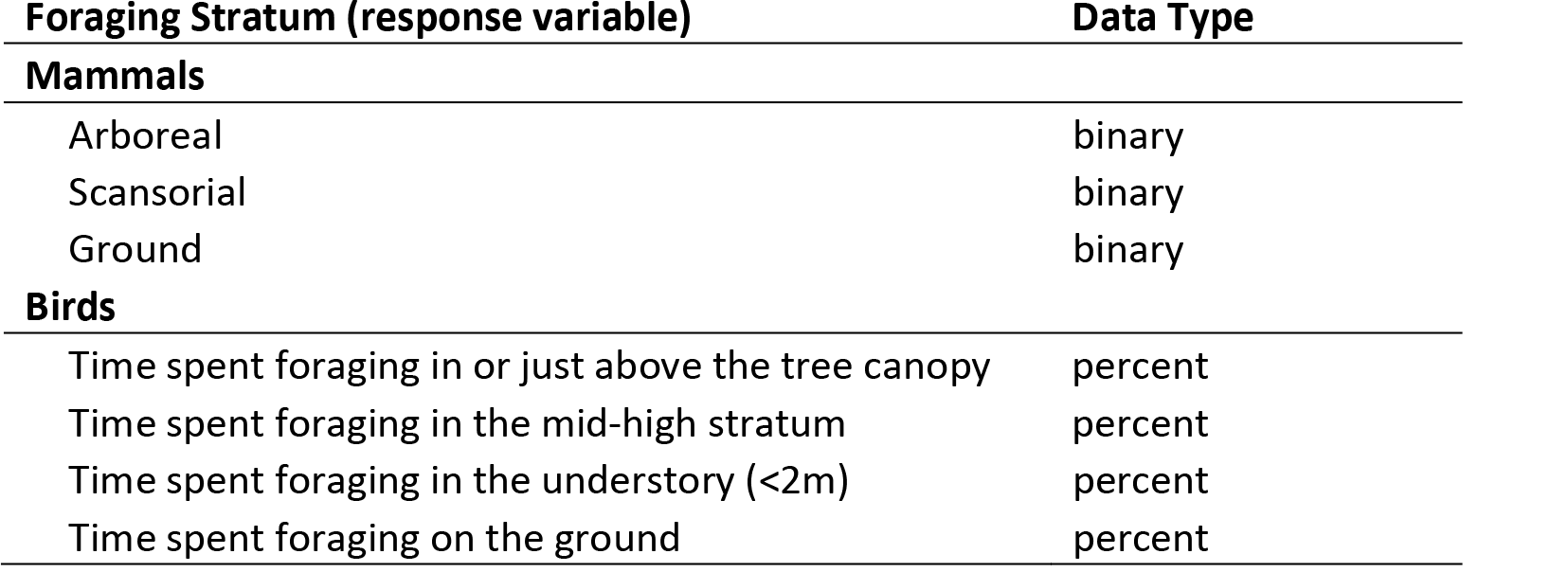
Foraging niches modeled for mammals and birds and their associated data types.

### Phylogeny

To account for shared evolutionary history, we acquired consensus phylogenetic trees for mammals (Faurby et al., 2018) and birds (Jetz et al., 2012) and used them to calculate phylogenetic distances, based on branch lengths, between each species and every other species. We conducted a principal components analysis on the phylogenetic distance matrix and extracted the first two components (Diniz-Filho et al., 2012; Santini et al., 2022). The first component, referred to here as Phylo1, generally represents larger phylogenetic distances between clades closer to the root of the tree, while the second component, referred to here as Phylo2, generally represents smaller distances between more recently evolved groups (Diniz-Filho et al., 2012). The first two principal components explained 91% of variance for mammals and 72% for birds.

### Predation

To estimate the influence of terrestrial and aerial predation on mammal and bird vertical foraging niche, we linked each species in the EltonTraits database to its International Union for the Conservation of Nature (IUCN) and BirdLife International and Handbook of the Birds of the World (BOW) geographic range (BirdLife International and Handbook of the Birds of the World 2018, IUCN 2019). We defined terrestrial and aerial predators as animals foraging on the ground and in the air whose diet is comprised of at least 25% vertebrate endotherms. Additionally, many large predators have recently gone extinct. The Phylacine database contains information on diet, body mass, and natural range of extinct mammals (Faurby et al., 2018). We used this database to identify extinct terrestrial predators whose diet is comprised of at least 25% vertebrate endotherms. The database does not contain information on foraging stratum of extinct mammals but examination of those species with diets comprised of at least 25% endotherms suggests that most of them had primarily ground foraging habits. We removed two species that appeared to be primarily marine foraging, *Neomonachus tropicalis* and *Zalophus japonicus*. We conducted sensitivity tests on the vertebrate endotherm threshold for defining predators, using alternative values of 10% and 50%. Results were similar for these thresholds so in the main text we report results using the 25% threshold.

For each species, we used predator-prey body size relationships (Carbone et al., 1999) to identify a candidate set of predators that could potentially consume that species. This candidate set was defined as predators with a probability of at least 0.01 of capturing prey of the given body mass according to equations that model prey body size as a roughly normal distribution around an optimum value (Harfoot et al., 2014; Hoeks et al., 2020). For smaller mammal predators, < 21 kg, optimal prey size is 10% of predator mass. For larger mammal predators, >= 21 kg, optimal prey size is equal to predator mass.

Information on predator-prey ratios for predatory birds is scarcer than for mammals. Based on published accounts, however, we assumed optimal prey size for smaller predatory birds, < 2 kg, is 10% of predator mass (Marti et al., 1993). For larger predatory birds, > 2 kg, we assumed optimal prey size is 40% of predator mass (Bedrosian et al., 2017; Ibañez et al., 2003; Marti et al., 1993; Tarboton and Allan, 1984). The 2 kg threshold differentiates eagles, large hawks, and eagle-owls from smaller hawks, owls, and falcons.

We then conducted a spatial overlay analysis to determine whether any of candidate predators had at least 10% range overlap with the focal species. For those predator species, we then used equations from Hoeks et al. (2020) to calculate the probability of capture for the focal species, ranging from 0.01 to 1 and summed them to create an indicator of potential predation pressure. To conduct range overlays, we reprojected IUCN and BOW range maps to the Behrman cylindrical equal area coordinate reference system and converted them to raster format with a 20 km cell size. This is slightly coarser cell size than recent global analysis of species ranges using a 10 km cell size (Jenkins et al., 2013) but was chosen to facilitate computation while still capturing relevant details of species ranges. We reprojected and resampled Phylacine ranges to match the rasterized IUCN and BOW ranges.

Before modeling and mapping (see below), we filtered the data for species that were present in the Elton traits databases, the IUCN and BOW range databases, and phylogenetic databases, resulting in 4828 out of 5400 mammal species and 9437 out of 10,203 bird species with data on traits, phylogenetic distance, and geographic ranges.

### Mapping the Vertical Foraging Niche

To provide geographic context to our analyses, we linked vertical foraging niche information to range maps of birds and mammals. We then converted each species range to raster format. For mammals, we summed the number of species in each vertical foraging niche in each cell and divided by the total number of mammal species in that cell. For birds, we summed the percent time foraging in each vertical niche for each species in a cell, then divided the total by 100. We then divided this by the total number of bird species in that cell.

### Predicting the Vertical Foraging Niche

We used the random forest machine learning algorithm (Breiman, 2001) to model foraging stratum in mammals and birds as a function of intrinsic functional traits such as diet and body mass, phylogeny, and extrinsic predation pressure (Table 2). Activity period may have direct relationships on the vertical niche as in the case of diurnal primates that use color to differentiate between high- and low-quality fruits. It may also have an indirect influence on the vertical niche via its relationship with other variables such as predation or body mass (Cox et al., 2023, 2021). For example, larger body size is thought to be a defense against predation while illumination is thought to increase predation risk. The relatively larger body size of most diurnal primates may be an adaptation to increased predation risk of daytime activity (Burnham et al., 2013). Before fitting random forest models, we calculated correlations between variables to identify multicollinearity that may affect interpretation of variable importance values. We observed only two correlations >= ∼ 0.7, between passerine/non-passerine and Phylo1 variables for birds, and between nocturnal and diurnal variables for mammals. We omitted the diurnal variable from models for mammals and the passerine/non-passerine variable from models for birds, as Phylo1 was judged to contain more information relevant for understanding the vertical niche.

**Table 2.** Predictor variables used in regression models of foraging stratum for birds and mammals. Diet variables represent the percent of that food type in the diet of each species. See methods for explanation of variables developed in this study.

| Description | Units | Source |
| --- | --- | --- |
| Body mass | kg | EltonTraits |
| Diet<sup>†</sup> |  |  |
| fruit | % | EltonTraits |
| nectar | % | EltonTraits |
| seed | % | EltonTraits |
| plant (other) | % | EltonTraits |
| vertebrate (endotherm) | % | EltonTraits |
| vertebrate (ectotherm) | % | EltonTraits |
| fish | % | EltonTraits |
| unknown vertebrate | % | EltonTraits |
| scavenged | % | EltonTraits |
| Activity period |  |  |
| Nocturnal | Boolean | EltonTraits |
| Crepuscular* | Boolean | EltonTraits |
| Phylogenetic distance (Phylo1, Phylo2) | Unitless | This study |
| Predation pressure | Unitless | This study |

\*Not available for birds, <sup>†</sup>Provided in 10% increments

For both mammals and birds, we used randomized search with 5-fold cross-validation to test 100 candidate sets of random forest hyperparameters which included number of trees, maximum tree depth, maximum number of predictors to use, minimum number of samples required to split a node in a tree, minimum number of samples required to be in a leaf in a tree, and whether or not to use the whole dataset when building trees. Once the best candidate set of hyperparameters was identified, we fit a model using those hyperparameters on an 80/20 training-test split which we used to calculate AUC and F-scores for mammals and RMSE and r^2^ for birds. Different model performance metrics are required because the response variables for mammals are binary (e.g. arboreal and not-arboreal) while response variables for birds are percent (e.g. percent time foraging in the mid-high canopy). We then fit models using the whole dataset for assessing variable importance and mapping geographic distributions of variable effects. To better understand the influence of phylogeny on vertical niche as well as interrelationships with functional traits, we also fit models without phylogenetic variables Phylo1 and Phylo2. While the response variables for mammals are binary, random forests supports continuous predictions of the probability of being in a particular foraging niche, calculated as the proportion of trees in a random forest predicting membership in that foraging niche. We used the probability output format for mammals for calculating Shapley values (see below).

We assessed variable importance for predicting vertical foraging niche using Shapley values (Lundberg et al., 2020), a model-agnostic, game theoretic estimate of the additive contribution of each predictor variable to each prediction. In simpler terms, Shapley values quantify how much a variable increases or decreases a given prediction relative to the expected average prediction. In the case of a binary response variable, the average prediction is the proportion of the target class in the dataset (e.g. the proportion of arboreal mammals). In the case of a continuous response variable, the average prediction is the average value of the response variable (e.g. average % time spent foraging in the mid-high canopy). To calculate the Shapley value of a target variable *V* for a specific species, a prediction for that species is made using a given combination of the predictor variables without *V*. The difference between this prediction and the average prediction is calculated. Another prediction is then made but this time including *V*. The difference between these two predictions is the marginal contribution of *V*. The Shapley value for *V* is the average of the marginal contribution of *V* across all possible combinations of predictor variables for a specific species. The average absolute value of Shapley values for a given variable across all species can then be used as an indicator of that variable’s importance.

We used Shapley values to visualize and rank predictor variable importance and to create response curves for predictor variables to estimate their effects. Body mass was log-transformed for display in response curves. We did not transform body mass before model fitting as tree-based models are invariant to monotonic transformations of predictor variables (De’ath and Fabricius, 2000). However, we conducted tests where we log transformed body mass before model fitting to determine whether Shapley values were sensitive to transformation. Results from these models were virtually identical to our original models so we report on the original models in the main text. We also summarized Shapley values by taxonomic family to characterize the distribution of importance values across families.

Response curves and summaries by taxonomic family were developed only for the most important variables within and across foraging strata.

### Mapping Geographic Distributions of Vertical Foraging Niche-Predictor Relationships

To provide context for the analysis of vertical niches, we linked Shapley values for each species and predictor variable to IUCN and BOW range maps to visualize variable importance by foraging stratum for mammals and birds for only the most important variables. For a given predictor variable for either mammals or birds, we attributed the Shapley value to each species’ range and then summed overlapping ranges gridded at 20 km. We divided this sum by the total number of species in each grid cell, resulting in an average importance value for each grid cell for each variable for each stratum.

Throughout, we omitted cells covered by >= 50% water.

## Results

### Geographic Distributions of Mammal and Bird Foraging Niches

Areas of high prevalence of arboreal mammals were in the Caribbean, the Amazon Basin, West Africa, and throughout Southeast Asia (Figure 1). Scansorial mammals had high prevalence throughout much of tropical Central and South America, Western Europe, Madagascar, and mainland Southeast Asia. Areas of high prevalence of ground mammals were widely distributed in temperate, cold, and xeric regions.

**Figure 1.**
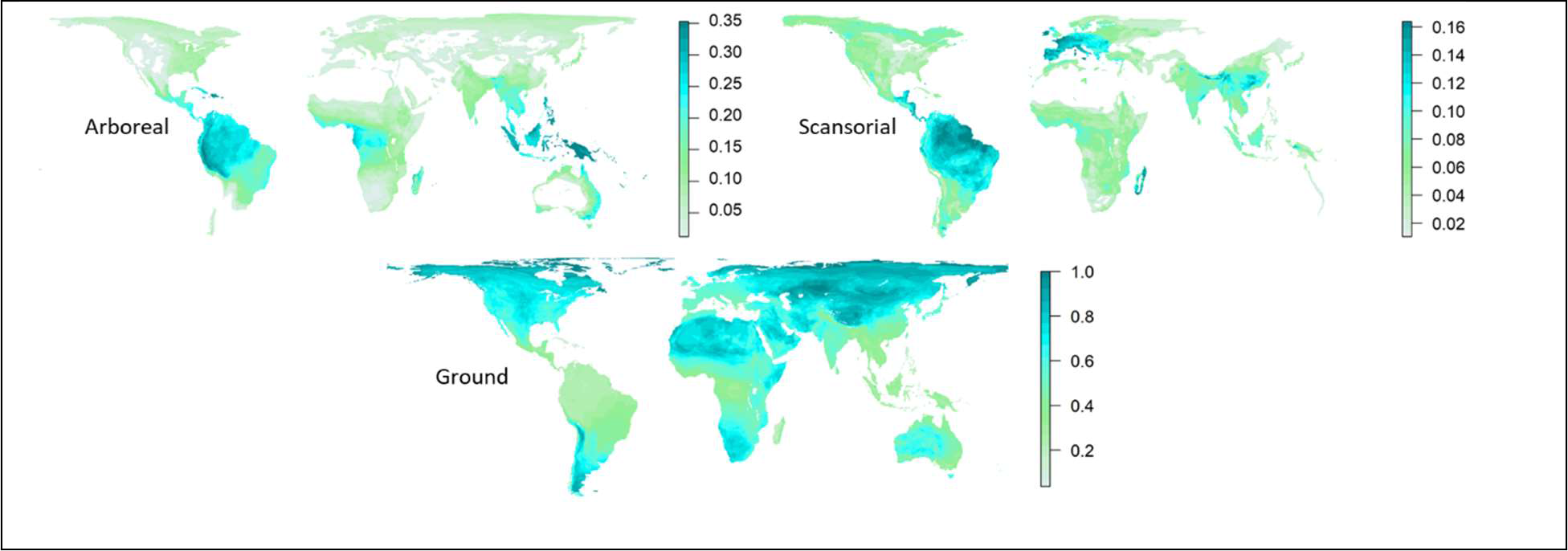
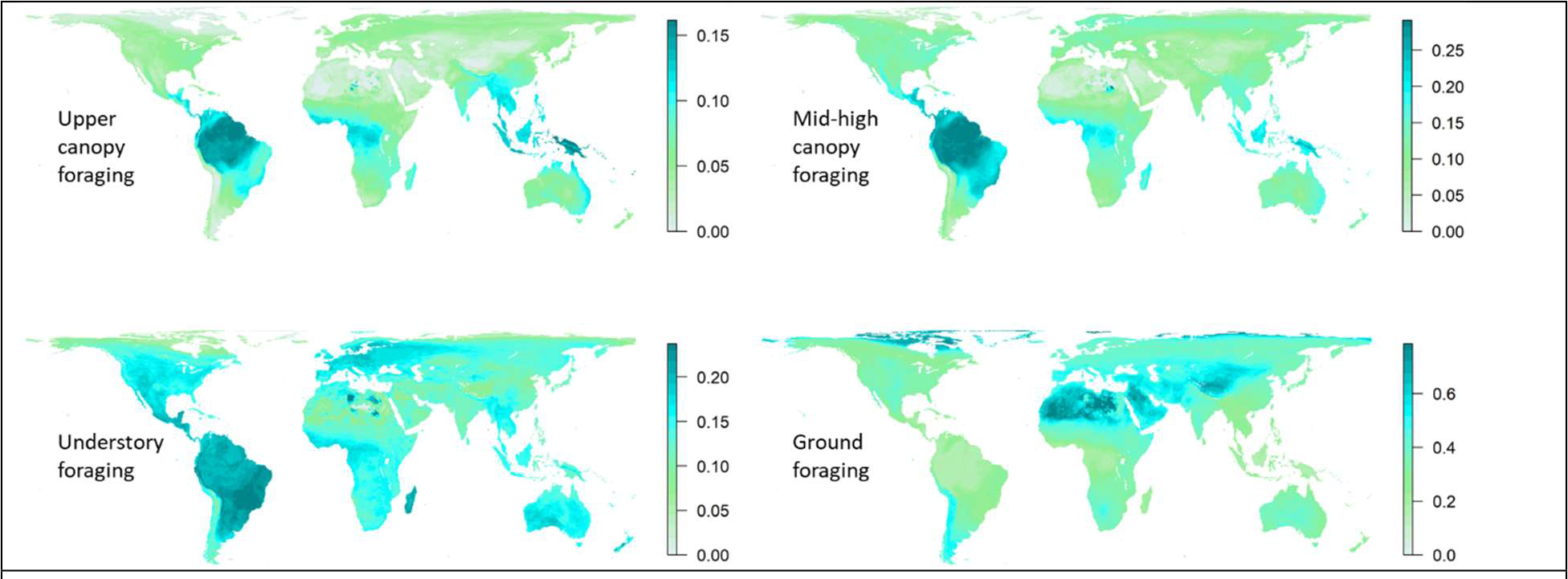
Richness of mammals in each vertical foraging niche, normalized by the total number of mammal species (top two rows). Sum of percent time birds spent foraging in each vertical niche, divided by 100, and normalized by the total number of bird species (bottom two rows).

For birds, areas of high prevalence for upper canopy foraging were distributed across the tropics with the highest concentrations in the Guiana Shield and the island of New Guinea. Areas of high prevalence for mid-high canopy foraging were similar to those for upper canopy foraging but with somewhat lower prevalence in Central Africa and Southeast Asia and somewhat higher prevalence in the Amazon Basin and the Atlantic Forest. Understory foraging was more widespread, with the highest prevalence in the Cerrado and Atlantic Forest in South America, Madagascar, and isolated locations in Northern Africa.

Ground foraging was concentrated in cold and xeric regions with the highest concentrations in Northern Africa, the Middle East, the Tibetan Plateau, and the Arctic.

### Predictors of Mammal Vertical Foraging Niche

Models predicting mammal vertical foraging niche performed well, with AUC scores above 0.96 for all strata and F1 scores above 0.95 for arboreal and ground strata and above 0.78 for the scansorial stratum. In terms of overall importance, fruit and invertebrate consumption were the strongest predictors of arboreal foraging (Figure 2). Both Phylo1 and Phylo2 were important for predicting all foraging niches.

**Figure 2.**
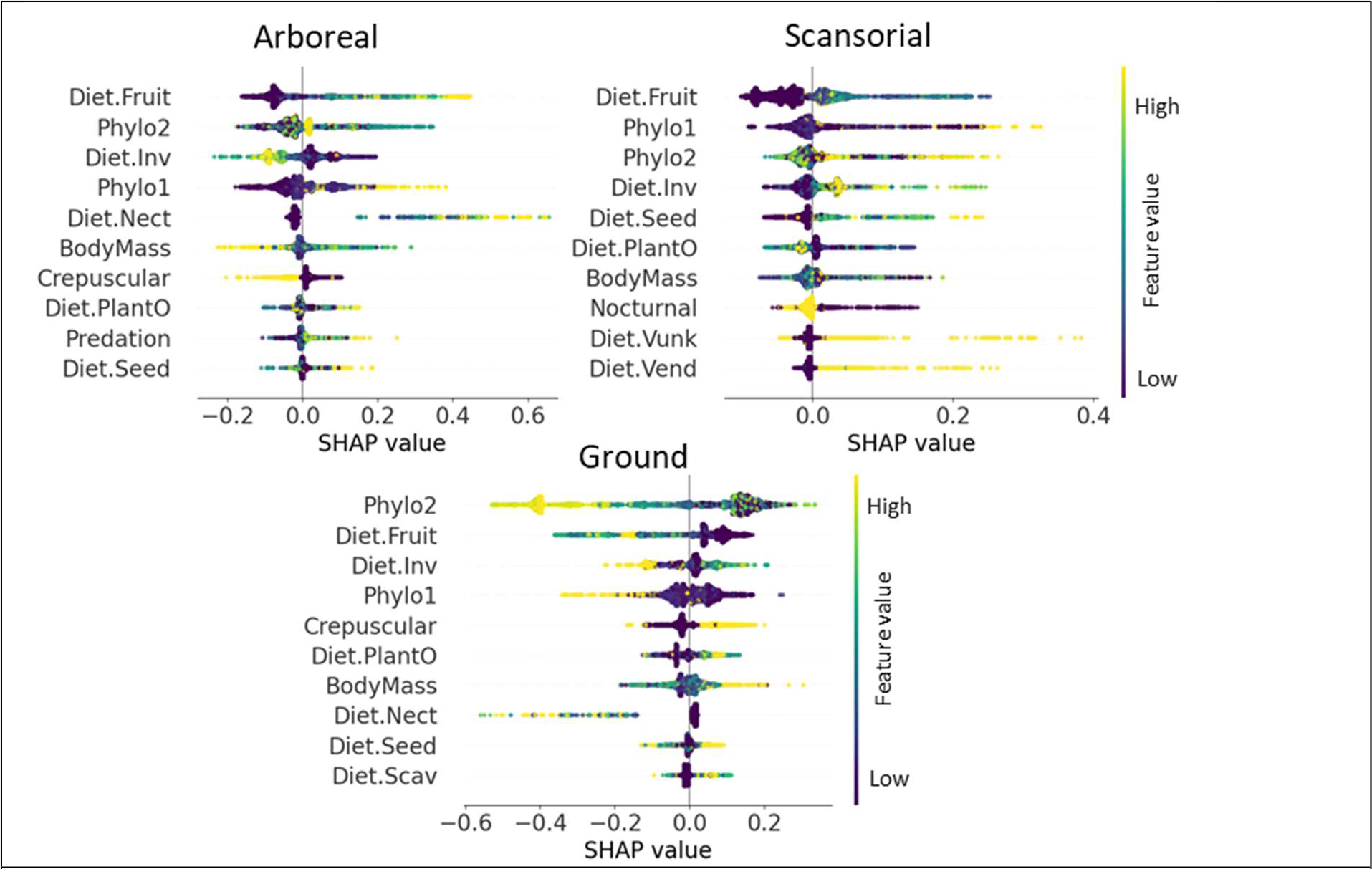
Shapley values (SHAP) for arboreal (top left), scansorial (top right), and ground (bottom) foraging mammals. Each point represents a species. The yellow to purple color ramp represents high to low values respectively, of the predictor variables. Variables with a “Diet” prefix indicate percent diet composed of a given food type. Inv=invertebrates, Scav=scavenged, Nect=nectar, PlantO=other plant material (not fruit, nectar, or seeds), Vect=vertebrate ectotherms, Vend=vertebrate endotherms, VFish=vertebrate fish, Vunk=vertebrate unknown. Phylo1 and Phylo2 are the first two principal components of the phylogenetic distance matrix. Variables are ordered by the mean absolute value of Shapley values. Note different x-axis scales. For clarity, only the top ten variables are shown.

Fruit and nectar consumption were both strongly positively related to arboreality while invertebrate consumption was negatively related. Nectar consumption was strongly and positively important for arboreal foraging for some species but for most its influence was near zero. Small to medium body mass values (0.5 – 12 kg) were positively related with arboreality while crepuscular activity was negatively related.

Strong functional trait predictors for scansorial mammals were more varied and included consumption of fruit, invertebrates and seeds. Body mass was important but its influence was variable with positive peaks at ∼5 g, ∼80 g, and ∼5 kg and a negative peak around 20 g. As with arboreal mammals, both Phylo1 and Phylo2 were highly important. % vertebrates in diet had minimal influence for most species but for those where it was important, such as for scansorial consumption of unknown vertebrates or endotherm vertebrates, it was strongly positive.

Phylo2 was most important for predicting ground foraging in mammals. Fruit consumption was strongly negatively correlated with ground foraging. Invertebrate and plant consumption were important, with medium values positively associated with ground foraging but with high and low values negatively associated with it. Very high body mass values were positively associated with ground foraging while medium and low body mass values exhibited variable influence although the overall importance of body mass was low. Predation was relatively unimportant across strata although it was among the top 10 most important variables for arboreal foraging.

### Predictors of Bird Vertical Foraging Niche

Models predicting vertical foraging niche for birds performed moderately, with r^2^ ranging from 0.28 for understory birds to 0.53 for ground birds. RMSEs ranged from 19.53 for canopy birds to 26.14 for ground birds. Fruit consumption was the most important predictor for upper canopy and mid-high canopy foraging birds and was positively associated with those niches in both cases (Figure 3). Phylo1 and body mass were the next most important variables, respectively, for upper canopy and mid-high canopy foraging birds. Body mass was strongly negatively associated with upper canopy and mid-high canopy foraging. As with arboreal mammals, nectar consumption was not important for most species, evidenced by the large cluster of points near the zero line. For those which it was, it was positively and strongly associated. A similar but reversed pattern was seen for seed consumption in mid-high canopy foraging birds where it was unimportant for most species but had a strong negative influence for some.

**Figure 3.**
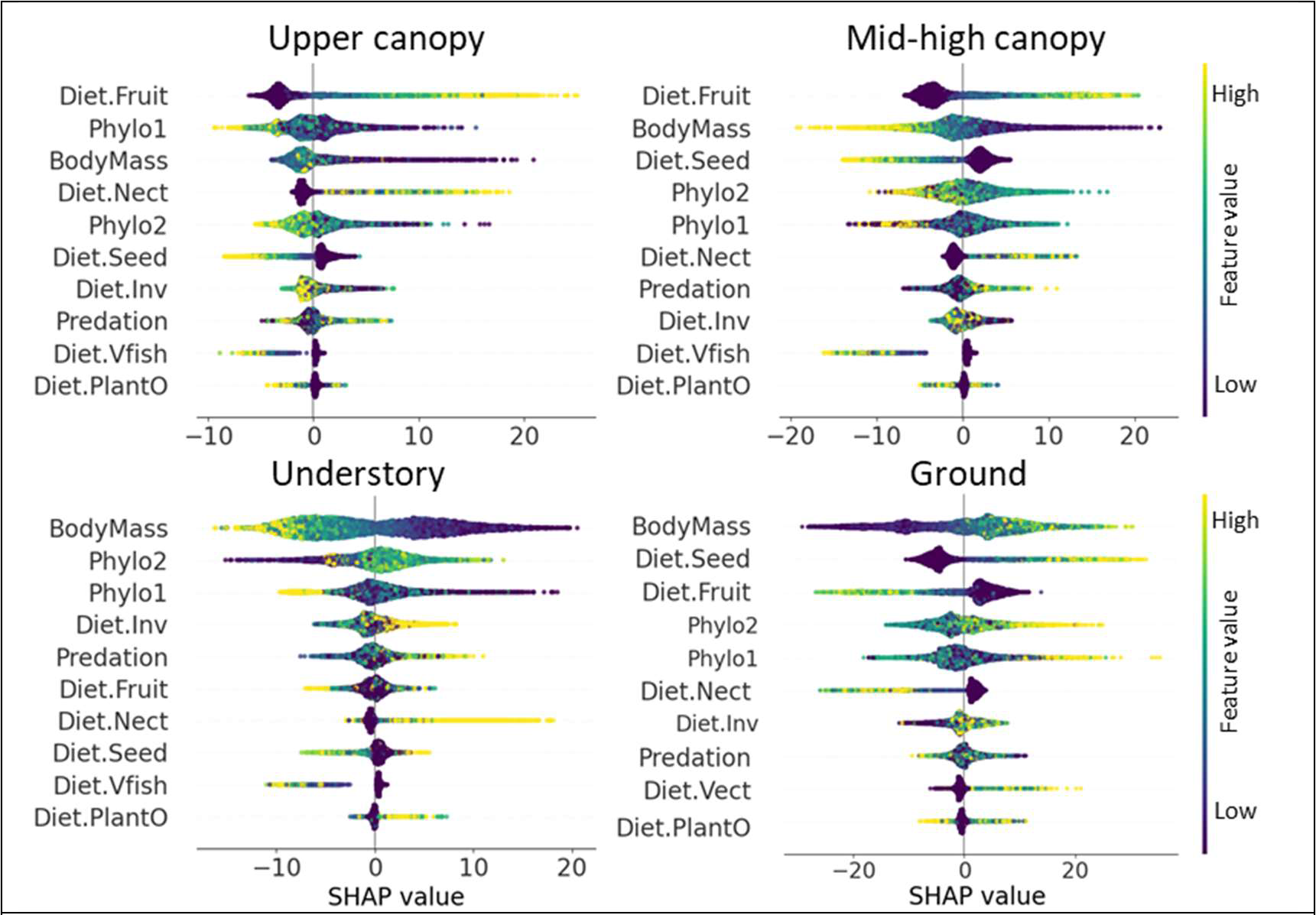
Shapley values for upper canopy (top left), mid-high canopy (top right), understory (bottom left) and ground (bottom right) foraging birds. Each point represents a species. The yellow to purple color ramp represents high and low values, respectively, of the predictor variables. Variables with a “Diet” prefix indicate percent diet composed of a given food type. Inv=invertebrates, Scav=scavenged, Nect=nectar, PlantO=other plant material (not fruit, nectar, or seeds), Vect=vertebrate ectotherms, Vend=vertebrate endotherms, VFish=vertebrate fish, Vunk=vertebrate unknown. Phylo1 and Phylo2 are the first two principal components of the phylogenetic distance matrix. Variables are ordered by the mean absolute value of Shapley values. Note different x-axis scales. For clarity, only the top ten variables are shown.

For understory and ground foraging birds, body mass was the most important variable but had opposite direction of influence in each. Body mass was strongly negatively associated with understory foraging while it was strongly positively associated with ground foraging. Phylogenetic variables were the next most important for predicting understory foraging. Seed and fruit consumption were important for predicting ground foraging with strongly positive and negative relationships, respectively. Importance values for predation were generally small for most species in all strata but for species where it was important, its influence was variable.

### Cross-Stratum Comparison of Mammal Predictors

Fruit and invertebrate consumption were the most important functional trait predictors for vertical foraging niche in mammals and their partial response curves showed systematic variability across strata (Figure 4). The influence of fruit consumption became progressively less positive when moving from arboreal to ground foraging strata. For arboreal mammals, the importance of fruit consumption exhibited a strong positive trend. For scansorial mammals, the relationship was unimodal with a peak at around 30% of fruit in diet. For ground foraging mammals, the relationship exhibited a negative trend from 0-∼30% fruit consumption before leveling off at negative values. Invertebrate consumption in arboreal mammals exhibited a largely negative trend from 0-60% consumption before leveling off at negative values. In contrast, trends for scansorial and ground foraging mammals were weakly positive from ∼25-60% invertebrate consumption whereafter the influence of invertebrate consumption was largely positive. The primary exception to this trend was in ground foraging mammals with 100% invertebrate consumption. For these species the relationship between invertebrate consumption and ground foraging was generally negative. Predation was relatively unimportant for scansorial and ground foraging mammals but displayed a small positive trend for arboreal foraging species, with most Shapley values above zero at an average predation value of 0.5.

**Figure 4.**
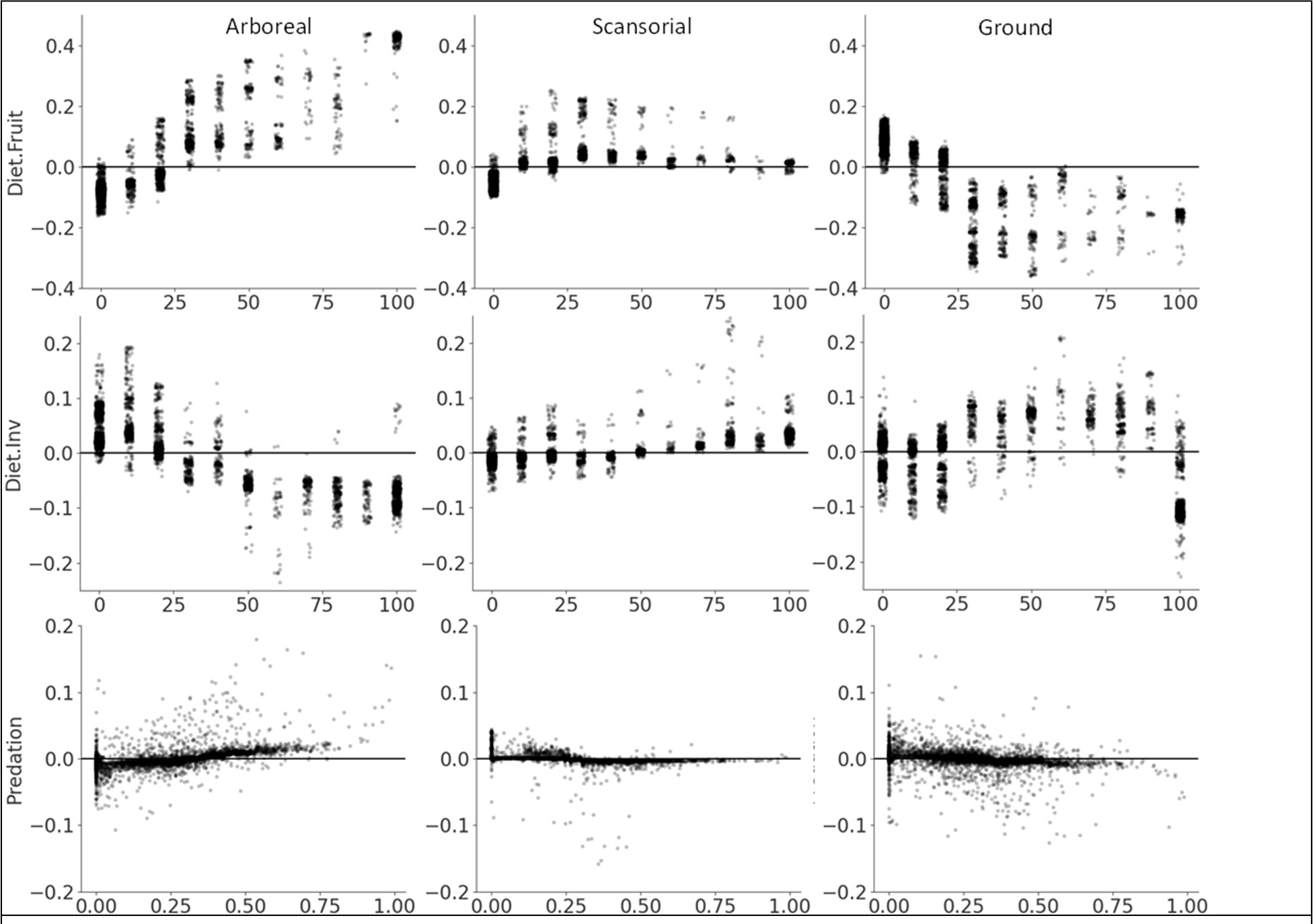
Response curves by vertical foraging niche for the two most important function trait predictors for mammals (top two rows) and predation (bottom row). Each point represents a species. Predictor variables are plotted on the x-axes. Shapley values are displayed on the y-axes and represent the marginal effect that each variable has on the predicted probability of a species being in each vertical foraging niche.

Phylo1 and Phylo2, indicators of shared evolutionary history derived from phylogenetic distance matrices, were frequently among the top three variables when predicting both mammal and bird vertical foraging niches. Similar values among species indicate similar evolutionary distances between those species and other species in the phylogeny. The interpretation of Phylo1 and Phylo2 is not as straightforward as for functional traits but summarizing their importance values by taxonomic order and family gives an indication of how they inform predictions of foraging niche (Figure 5).

**Figure 5.**
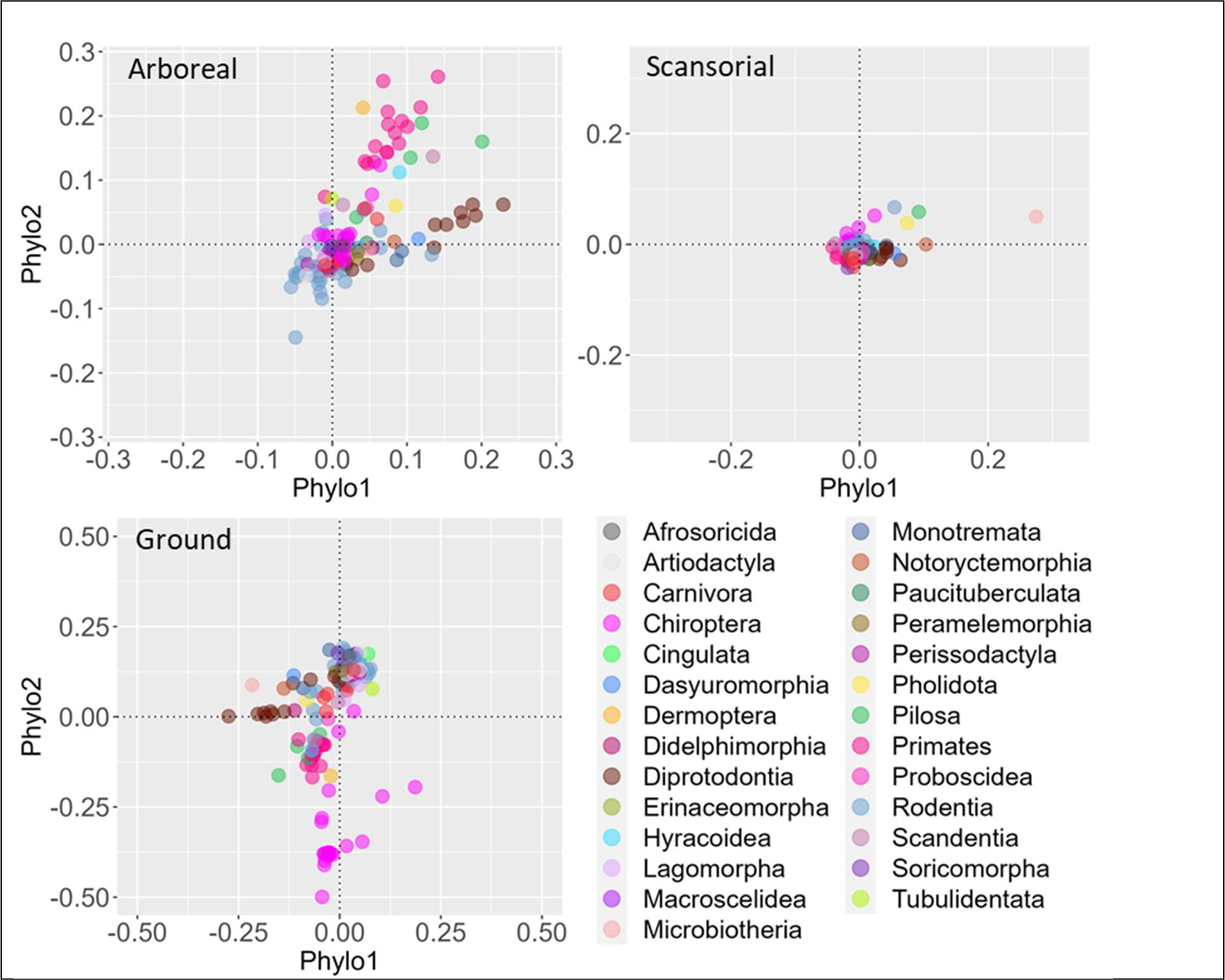
Scatterplot of Shapley values of Phylo1 and Phylo2 phylogenetic variables summarized by median value for taxonomic family and grouped by taxonomic order for arboreal, scansorial, and ground foraging mammals. Note different axis scales.

For the mammal arboreal niche, several families in the order Diprotodontia (marsupials) had the highest positive values for Phylo1 (larger phylogenetic distances associated with root clades), as did Dasyuromorphia (carnivorous marsupials) and one family of rodents. Several families in the order Primates had high positive values for Phylo2 (smaller phylogenetic distances associated with more recently evolved groups). Several smaller orders including Pilosa (sloths and anteaters), Dermoptera (gliding colugos), Hyracoidea (hyraxes), and Scandetia (treeshrews) also had high Phylo2 values. Most families of rodents had negative values for both variables.

For the scansorial mammal niche, most of the variability was on the positive side of the Phylo1 axis (note different axis ranges between Phylo1 and Phylo2). The highest values were seen in Microbiotheria (a small marsupial order) and Notoryctemorphia (marsupial moles). All values were quite small for Phylo2, ranging from ∼ -0.05 to 0.075, with a small number of families from Rodentia, Chiroptera (bats), Pilosa, Microbiotheria and Pholidota (pangolins) showing the highest values.

For the ground mammal niche, most of the variability was on the negative side of Phylo1 and Phylo2 axes, primarily due to negative values for several Diprotodontia and Chiroptera families, respectively. Phylo2 also had negative values for several families in Primate and Pilosa orders. Most of the other families had smaller positive values for Phylo1 and Phylo2, especially families in Rodentia.

### Cross Stratum Comparison of Bird Predictors

Body mass and percent fruit in diet were the top functional trait predictors for birds and their importance varied systematically across strata (Figure 6). Body mass exhibited a negative trend for upper canopy foraging birds which became progressively more negative for mid-high and understory foraging birds. However, ground foraging birds exhibited a strong positive relationship with body mass. Fruit consumption displayed a strong positive trend for upper canopy foraging birds, a moderate positive trend for mid-high canopy foraging birds, no trend for understory birds, and a moderate negative trend for ground foraging birds. Predation pressure displayed a noisy relationship with all bird foraging niches, with numerous species responding either positively or negatively at most levels of predation. However, the response to predation was mostly positive for species in the mid-high canopy and understory foraging niches above an average predation pressure of 0.8.

**Figure 6.**
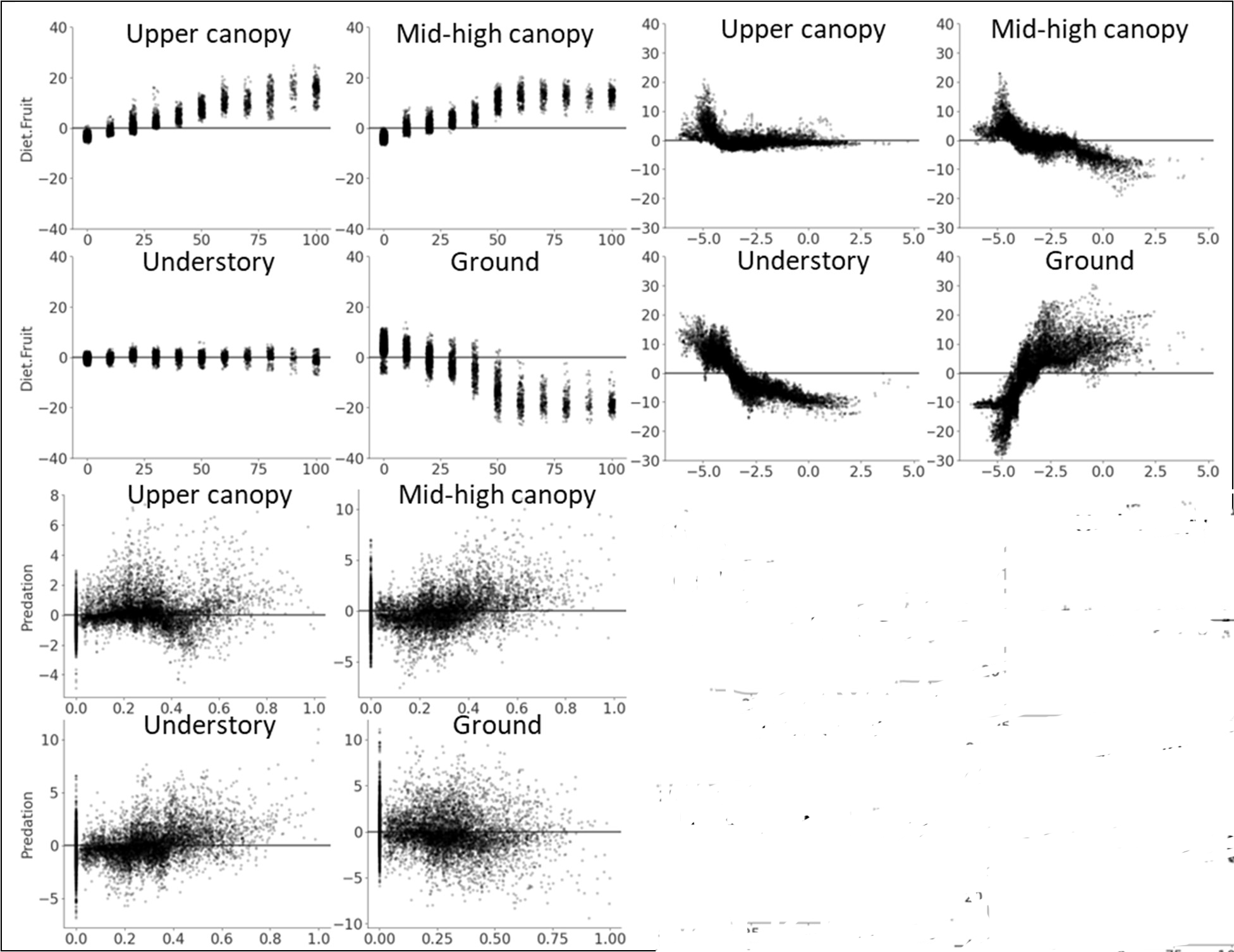
Response curves for the top performing functional trait variables - percent fruit in diet (top left panels), body mass (top right panels), and predation (bottom left panels) by foraging stratum for birds. Body mass values are log-transformed. Each point represents a species. Predictor variables are plotted on the x-axes. Shapley values are displayed on the y-axes and represent the marginal effect that each variable has on the predicted probability of a species being in each vertical foraging niche.

For the upper canopy bird foraging niche, families in Passeriformes had several families with both high positive and low negative values for both Phylo1 and Phylo2 (Figure 7). Families in Picciformes also had relatively high values in either Phylo1 or Phylo2Families in Casuariiformes, Gruiiformes, and Rheiformes had the most negative values for Phylo1.

**Figure 7.**
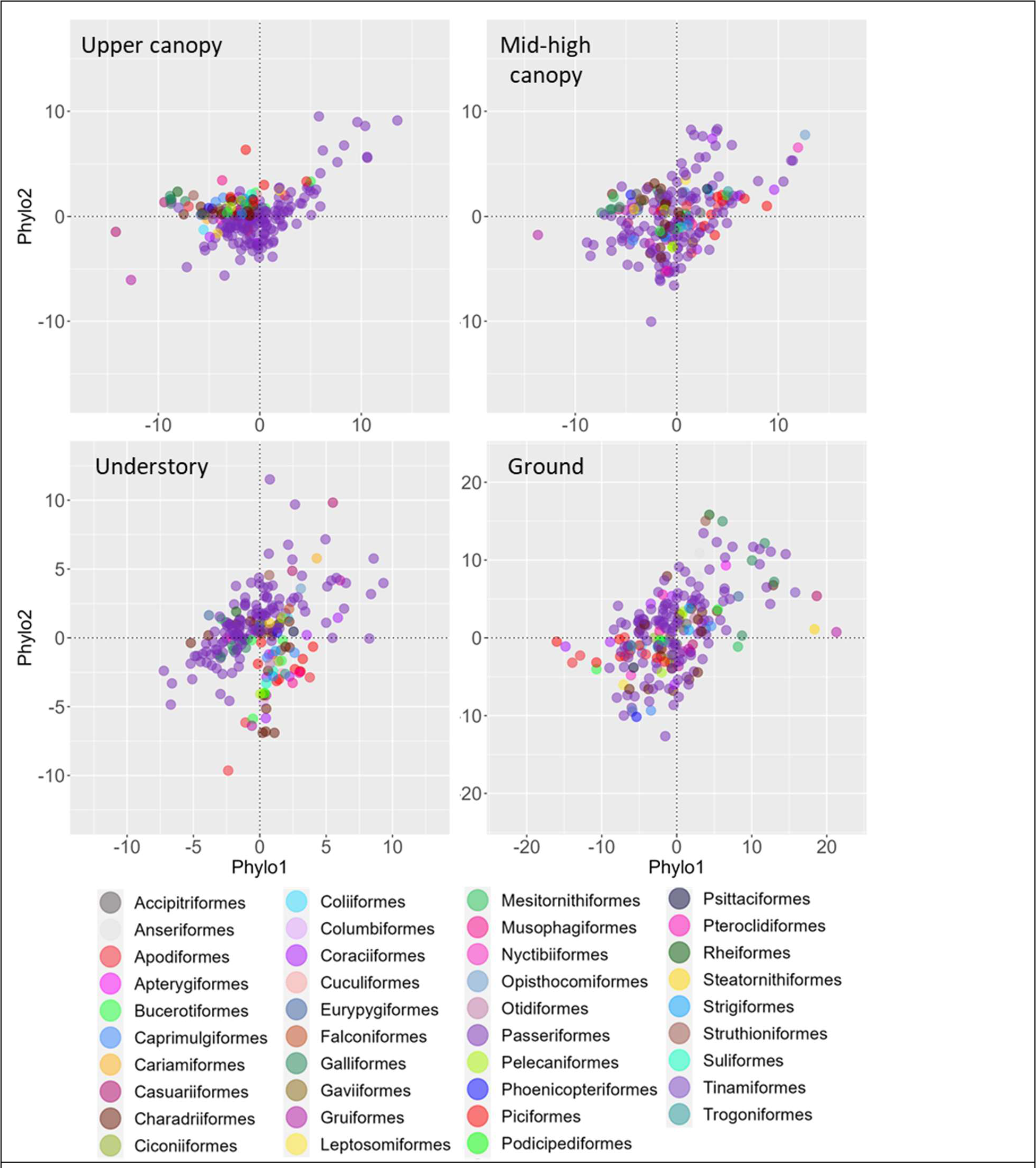
Shapley values of PC1 and PC2 phylogenetic distance variables summarized by median value of taxonomic family for bird foraging niche. Note different axis scale for ground foraging birds. Points are colored by taxonomic order and correspond to the legend labels.

As with the upper canopy niche, several families in Passeriformes had both high positive and low negative values for Phylo1 and Phylo2 for the mid-high foraging niche. Families in Opisthocomiformes, Nyctibiiformes, Coraciiformes, Piciformes also had high values for Phylo1. One family in Gruiiformes had the lowest negative value for Phylo1.

In the understory foraging niche, several families in Passeriformes had the highest positive values for both Phylo1 and Phylo2 and the lowest negative values for Phylo1. A small number of families in <u>Casuariiformes</u>, Cariamiformes, and Coraciiformes also had relatively high positive values for Phylo1 and Phylo2. The lowest values for Phylo2 were seen in several different families including Picciformes, Charadriiformes, Bucerotiformes, Coraciiformes, and Columbiformes.

### Effect of Omitting Phylogenetic Variables on Model Performance

Model performance for mammals was not substantially affected by omitting phylogenetic variables (difference of 0.07 at most) although performance on positive predictions for arboreal mammals, indicated by the F1-score, was slightly worse, 0.78 vs 0.72. Variable importance rankings and individual variable response curves were similar to those from models with phylogenetic variables included. Model performance for birds, however, declined moderately when phylogenetic variables were omitted, with a maximum decrease in r^2^ for ground birds of 0.16. Variable importance rankings were similar to models including phylogenetic variables. The response curves were similar as well, although omitting phylogenetic variables increased scatter.

### Geographic Patterns of Fruit Importance for Mammals

Fruit was the most important variable for predicting vertical foraging niche in mammals. Positive relationships between arboreal foraging and fruit consumption were overwhelmingly observed in the tropics with negative relationships virtually everywhere else. Hotspots of positive relationships included the tropical Andes, the coastal Afrotropics, and low land forests of the island of New Guinea. In the Indo-Australian archipelago, the island of Borneo marks a transition where Borneo and islands to the east show moderate importance of fruit while islands to the west show high importance of fruit for arboreal foraging. Positive relationships between fruit in diet and scansorial foraging were less prevalent than for arboreal foraging and were concentrated in the northern Amazon and Guiana shield in South America.

Very small areas of positive relationships were observed in the coastal Afrotropics, the eastern Himalaya, western New Guinea, and in the northern Nearctic. The pattern of fruit importance for ground foraging mammals was essentially the inverse of the pattern for arboreal foraging mammals. The strongest positive relationships were in continental interiors in North America, northern Africa, central and eastern Asia, and Australia. Other areas with positive relationships were the southern Andes, the Horn of Africa, and the northern Palearctic and Nearctic.

### Geographic Patterns of Fruit and Body Mass Importance for Birds

Fruit consumption was the most important variable for both upper canopy and mid-high canopy foraging birds. Positive relationships were observed almost exclusively in the tropics (Figure 8). The largest hotspots of positive relationships were in the tropical Andes, the Amazon, and lowland forests of New Guinea. Patterns for mid-high canopy foraging birds were largely similar with the strongest divergence in New Guinea. Here, the strongest positive relationships were in the central mountains rather than in lowland forests. Body mass was the most important variable for understory and ground foraging birds. Hotspots in positive relationships were sparse and concentrated in Central America and the tropical Andes. Smaller areas showing strong positive relationships included southeastern Brazil, north-central Africa, and the southern Arabian Peninsula. Positive relationships between body mass and ground foraging were widespread except for Central America and the Amazon where extensive negative relationships were observed. Hotspots of positive relationships included the eastern Sahara and northern portions of the Palearctic and Nearctic.

**Figure 8.**
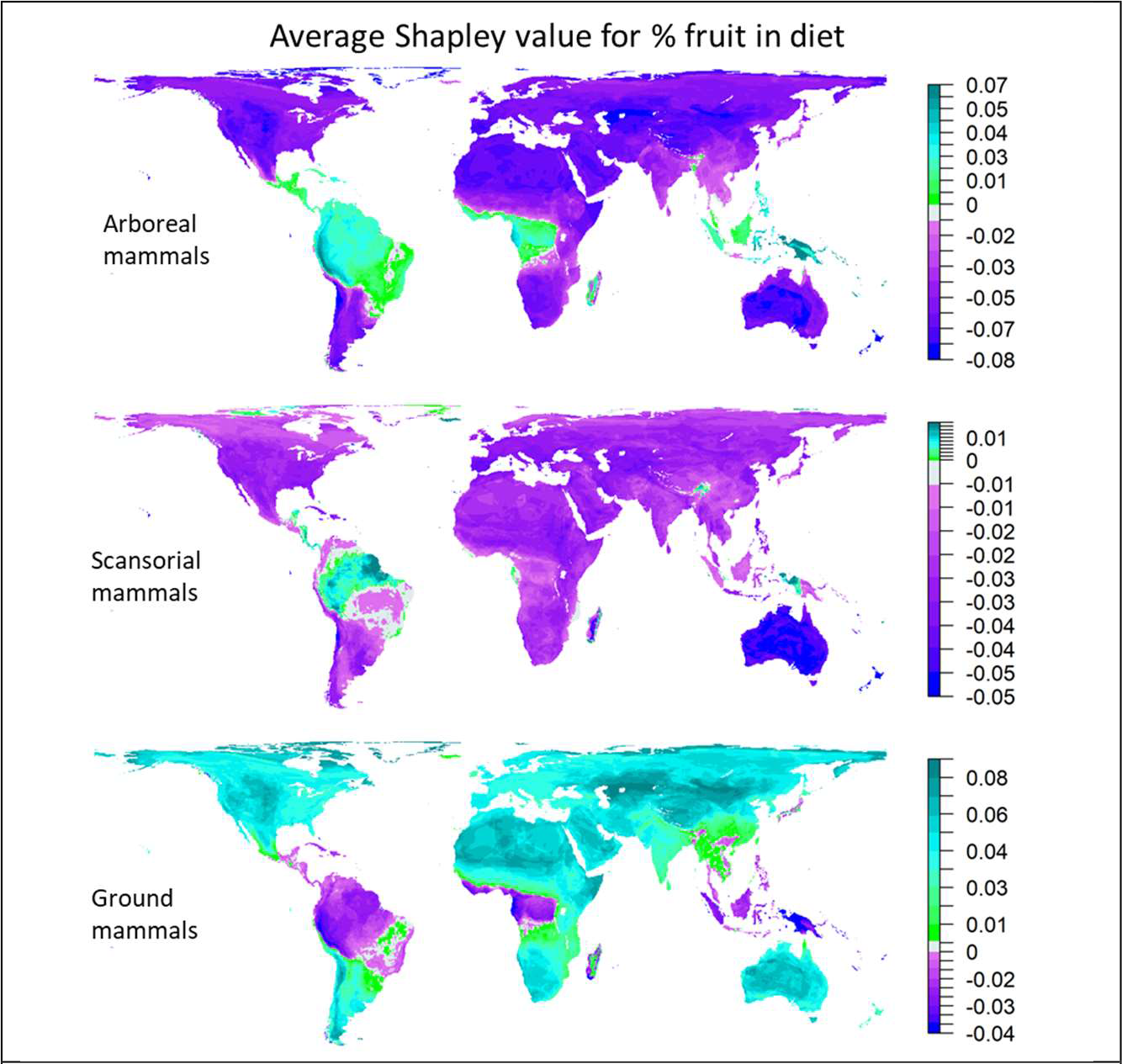
Average Shapley importance value for percent fruit diet for mammals.

**Figure 9.**
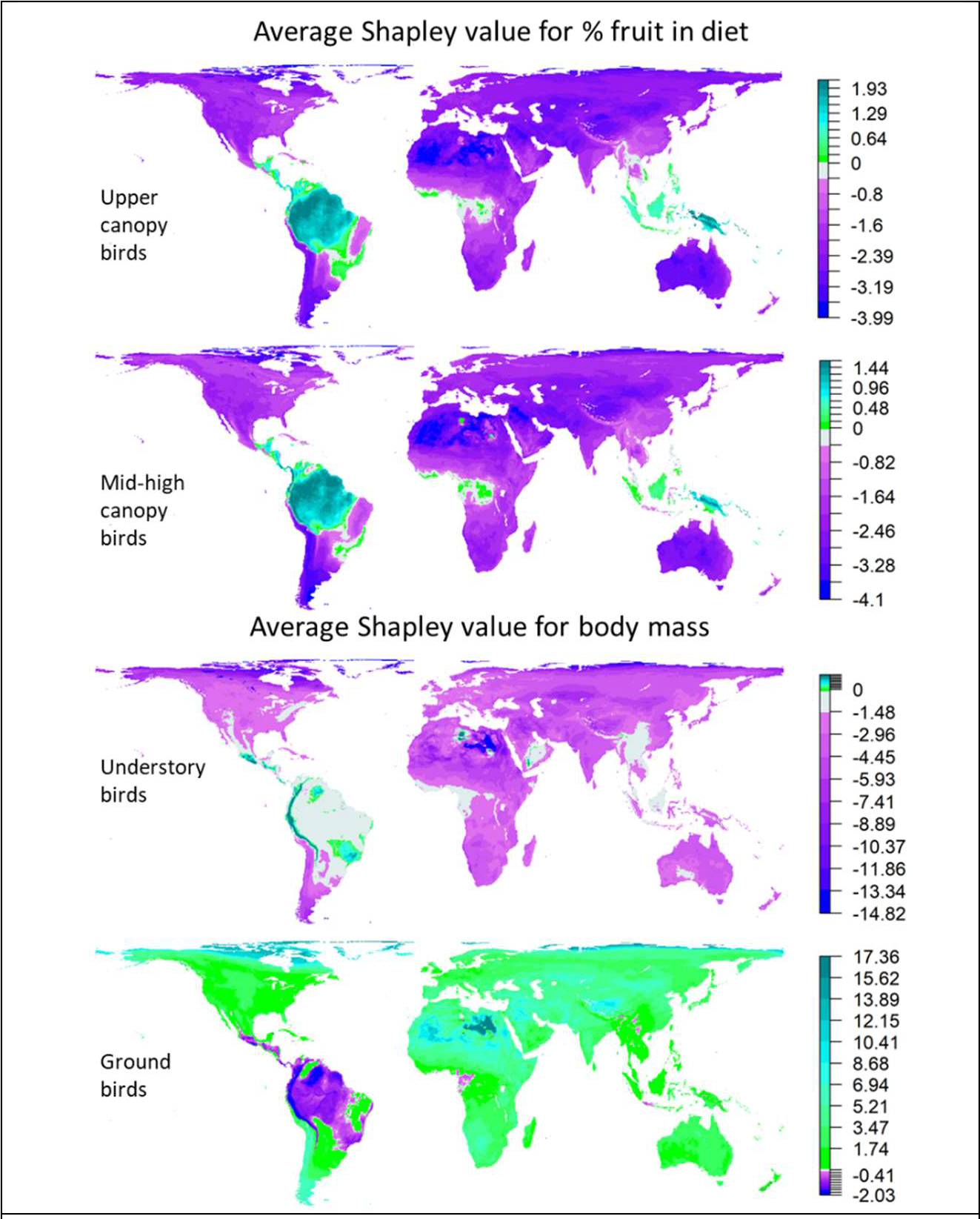
Average Shapley importance value for percent fruit in diet (top two panels) and body mass (bottom two panels) for birds.

## Discussion

Linking trait databases with species range maps revealed distinct global distributions of vertical foraging niches for mammals and birds (Figure 1). The most important predictors of these niches varied somewhat by vertical foraging niche and taxon but there were several systematic relationships. Diet was a strong predictor of vertical foraging niche across mammal and bird species. The distribution of canopy resources as a driving force in niche partitioning and community structure is a long-standing hypothesis with minimal evidential support at the macroecological scale (Fleming et al., 1987; Sussman et al., 2013; Tiffney, 2004). Our results support the idea of resource driven niche partitioning by showing that percent fruit in diet was positively associated with arboreal mammal foraging and upper and mid-high canopy bird foraging and negatively associated with ground foraging in both mammals and birds. While the thematic resolution of foraging strata in mammals and birds was limited to a handful of categories, models for both groups revealed progressively more positive relationships between frugivory and higher canopy foraging positions. This pattern could be explained by vertical resource gradients, and field observations have reported higher abundance of fruits and frugivores near the top of the canopy (Chmel et al., 2016; Francis, 1994; Houle et al., 2014; Pereira et al., 2010; Schaefer et al., 2002; Shanahan and Compton, 2001).

Consumption of invertebrates was also an important dietary predictor of mammal foraging niche which was positively associated with scansorial and ground foraging, displaying a negative relationship with arboreal foraging. This pattern may also be related to vertical resource gradients. Field inventories suggest invertebrate abundance is highest on or near the ground (de Souza Amorim et al., 2022; McCaig et al., 2020; Roisin et al., 2006) although Basset et al. (2015) found comparable numbers of arthropods in the upper canopy when compared to the ground. Perhaps more importantly, invertebrates on the ground are concentrated in the leaf litter and soil, facilitating easier capture than invertebrates distributed throughout the canopy. The negative relationship between high amounts of invertebrate consumption (∼100%) and ground foraging in mammals is likely attributable to bats which have several families with negative median importance values for the relationship between invertebrate consumption and ground foraging.

Body mass was not highly important for predicting mammal vertical foraging niche. This may be because phylogenetic variables are correlated with body mass, as was found with carnivores globally (Diniz-Filho et al., 2012), and therefore preempt body mass in the models. Models run without phylogenetic variables did result in an increase in the relative importance of body mass for scansorial and ground foraging mammals although the effect was small. Although not as important as diet and phylogeny, the shape of the response curves for body mass supports the idea of body mass as a constraint on vertical niches (Pineda-Munoz et al., 2016). Body mass for arboreal mammals showed a unimodal distribution, with positive importance values appearing between ∼5 g and 15 kg with a peak at 1 kg. Ground foraging mammals, in contrast, had largely negative body mass importance values at 1 kg but values were exclusively positive above 15 kg. Even though large animals such as gorillas and leopards frequently climb trees, food items in the canopy are generally found near slender branch terminuses, limiting the range of body sizes compatible with a purely arboreal lifestyle (Cristoffer, 1987). Several other theories exist explaining reduced body size in arboreal mammals including the biomechanical benefits of increased stability and increased movement and access across variable branch types and sizes, among others. Body size is just one morphological variable that we considered in our study but its behavior in our models suggest that biomechanics is an important constraint on arboreal living. Other notable morphological adaptations and strategies not included in our study include gliding, such as that seen in SE Asian and Australian mammals, as well as prehensile tails in New World monkeys.

Body mass was, however, highly important for predicting bird vertical foraging niche. Small body size up to ∼20 g was positively related to canopy foraging but showed essentially no relationship at larger body sizes. The relationship with body size became progressively more negative from the mid-high canopy to the understory. It then switched to a strongly positive relationship for ground foraging birds. These patterns may be due to resource gradients, competition, as well as vegetation structure gradients.

Recent advances in canopy profiling suggest that plant area density, a measure of vegetation distribution that includes leaves and stems, is generally highest near the ground and in the understory, becoming sparser near the upper canopy (Marselis et al., 2020; Milodowski et al., 2021; Schneider et al., 2019). While plant area may be highest near the ground, leaf area appears to show a peak near the top of some forest canopies (Almeida et al., 2019; Pearson, 1975; Stark et al., 2012). Vertical distributions of leaf and stem material likely influence substrates used for perching and hunting, type and density of food resources, and success in obtaining those resources (Pearson, 1975; Winkler and Preleuthner, 2001). For example, denser vegetation may be easier to navigate with a smaller body size and may also confer an advantage in capturing invertebrates (Pearson, 1975). Adaptations in birds to vertical niches may also influence macroevolutionary processes. Canopy bird species had lower genetic divergence than understory species across landscapes of Amazonia and the Andes (Burney and Brumfield, 2009). This pattern was largely attributed to dispersal propensity, which is higher in canopy than understory birds and related to wing morphology. Recent evidence points to strong collinearity between movement distances and body size in birds (Hartfelder et al., 2020) and thus we might expect large body size in canopy foraging species. Our results neither support or refute these expectations, but rather suggest a neutral relationship between canopy foraging and larger body sizes. Analysis of interactions between diet, body mass, and vertical foraging niche as well as additional traits related to locomotion and foraging such as wing, beak, and tail morphology (Burney and Brumfield, 2009; Pigot et al., 2020) could shed additional light on drivers of the observed relationships and potential macroevolutionary implications. In addition to more detailed analysis of traits, recent spaceborne lidar measurements reveal significant geographic heterogeneity in canopy structure (Doughty et al., 2023), leading to new opportunities to understand how current and historical vegetation structure may have influenced macroecological patterns.

The importance of variables that account for shared evolutionary history indicate that there are additional factors not represented by functional traits or predation that influence vertical niche occupancy. Our results also suggest that vertical niches are conserved in many mammal and bird groups. There were few instances of families with low Phylo1 scores (i.e. a negative association with a particular vertical niche) also having high Phylo2 scores (i.e. a strong positive association with a particular niche). This may reflect constraints of the body plan imposed by root clades on the ability of more recently evolved groups to occupy different vertical niches. These results also reveal the radiation that Passeriformes has undergone, enabling species in that group to occupy a wide array of vertical niches.

Other orders, such as Primates, Diprotodontia, and Pilosa tend to be strongly associated with a particular vertical niche and have diversified within it.

In contrast to diet, body mass, and phylogeny, predation was a relatively weak predictor of vertical foraging niche in mammals and birds. Its most systematic influence was on arboreal mammals, where higher predation pressure, at or above 0.5, was associated with a higher probability of arboreal foraging, providing some support for the landscape of fear hypothesis. Additional information on predator density and predator effectiveness may improve our understanding of the effects of terrestrial predation on the vertical foraging niche. There may also be countervailing relationships that reduce the effectiveness of the canopy as a refuge from predation. For example, numerous snakes, which we did not consider here, prey on smaller arboreal mammals and canopy foraging birds (Corlett, 2011). Additional reasons why we may not see a clear influence of predation pressure at macroecological scales may be attributed to the large number of other ways that animals can avoid predation including diel partitioning, crypsis (camouflage), morphological defenses (plating, spines, quills), chemical defenses, antipredator behaviors (increased vigilance, mixed-species herding), and large body size. While not an important predictor as formulated in this study, predation may still interact with functional traits and foraging niche to drive evolutionary processes via, for example, reduced mortality, access to higher quality food (e.g. fruit in the canopy), and longer life spans (Shattuck and Williams, 2010; Sibly and Brown, 2007). It is interesting to note that while there is significant arboreality in Australasia and Oceania, predation pressure from mammals and birds is relatively low. For example, New Guinea never had land connections with regions that hosted mammals of the order Carnivora, and no large marsupial predators evolved there (Corlett, 2011). This suggests that the benefit of increased access to canopy resources in these ecosystems may have been the main driver of the evolution of arboreal behaviors.

Mapped patterns of the influence of percent of fruit in diet suggest considerable geographic variability in the influence of functional traits on vertical foraging niche. Some of this variability is likely due to evolutionary contingencies in the distribution and diversification of mammal and bird species, some of which may be related to differences in forest composition and structure between realms. Lianas, for example, are relatively abundant in Neotropical forests but are relatively scarce in Indomalayan and Australasian forests (Emmons and Gentry, 1983). Such differences have been hypothesized to drive development of morphological adaptations such as tail prehensility and membranes for gliding that may help animals move between tree crowns with different amounts and types of structural connectivity provided by lianas (Emmons and Gentry, 1983). In both mammals and birds, the positive influence of fruit consumption on foraging in the canopy was primarily a tropical phenomenon. Geographic patterns suggest multiple potential mechanisms explaining hotspots of this relationship. In Southeast Asia, primates, which have a strong phylogenetic signal for the arboreal niche, are absent from eastern Indonesia and Australasia (Heads, 2010), with medium bodied arboreal foraging niches occupied by marsupials, which have a slightly weaker phylogenetic signal for the arboreal niche. We speculate that this manifests as an area of elevated positive relationship with fruit consumption to the east of Borneo, approximately congruent with Huxley’s modification of Wallace’s line delineating major zoogeographic regions (White et al., 2021). We observed a similar pattern in upper canopy and mid-high canopy foraging birds although the zone of strong positive relationship with fruit consumption was shifted east. This is, however, in agreement with a recent analysis of bird and mammal species turnover suggesting that birds of the Philippines have stronger ties with Asian lineages than mammals do (White et al., 2021).

In South America, the tropical Andes appear as a hotspot of the relationship between fruit consumption and arboreal foraging in mammals and, to a lesser extent, upper and mid-high canopy foraging in birds. Positive relationships between low body mass and understory foraging in birds was also pronounced in that domain. The Andes are a well-known biogeographic barrier, containing significant environmental gradients and many recently diversified species groups, as well as older lineages with high levels of endemism (Rahbek et al., 2019). We speculate that this overlap of numerous animal families with high diversity and an abundance of angiosperms (Pérez-Escobar et al., 2022) may explain high levels of fruit importance for canopy foraging in the tropical Andes.

Ground foraging birds show the strongest positive body mass relationships in areas of high environmental harshness, including the Sahara, Arabian Peninsula, continental Asia, and the far north. Herbivory in birds is relatively rare (e.g. Ptarmigans) but positive relationships between herbivory and body mass have been observed in several bird groups including large flightless and flight limited bird species like ostriches and emus (Olsen, 2015). Hotspots of high positive body mass relationships may reflect areas where herbaceous forage is more prevalent relative to other food sources and overall species numbers are low, allowing positive ground foraging body mass relationships to dominate. Positive body mass-ground foraging relationships at high latitudes may reflect Bergmann’s rule, where larger bodied animals are found in more poleward environments.

## Conclusion

We found strong relationships between functional traits, phylogeny, and vertical foraging niches of terrestrial mammals and birds, providing support for the theory of niche differentiation across vertical resource gradients at macroecological scales as well as support for vertical niche conservatism in many mammal and bird groups. Of the functional traits investigated, fruit and body mass were most important across taxa and vertical foraging strata. We found these functional traits were important even in the presence of strong phylogenetic signals for specific vertical niches in certain families and orders such as primates. When combined with the importance of fruit consumption for canopy foraging across multiple disparate families, these findings indicate convergent ecological adaptation to canopy niche spaces (Pigot et al., 2020). Geographic patterns in variable importance values suggest multiple mechanisms for spatial structure in eco-evolutionary relationships, including latitudinal gradients in vegetation structure and composition, historical patterns of island isolation (in Southeast Asia), and the influence of habitat heterogeneity driven by tectonic processes (in South America).

Moreover, our analysis of relationships between functional traits and foraging strata highlights potential impacts of climate change, deforestation, and forest degradation on species with particular functional traits. Climate change has induced reductions in flowering and fruiting of trees in Gabon, Africa (Bush et al., 2020). Large areas of the tropics are now dominated by structurally degraded forests which are shorter and structurally less complex than undisturbed tropical forest (Chazdon et al., 2016; Hansen et al., 2019). Climate change, forest loss, and forest degradation are thus likely to disproportionately impact animal communities that are adapted to canopy foraging (Laurance and Laurance, 1996; Whitworth et al., 2019). Loss of these animal species may in turn reduce seed dispersal and establishment of tree species that are characteristic of undisturbed forest, leading to shifts towards early successional, abiotically dispersed species at the landscape scale and ultimately a loss of functional diversity for multiple taxa (Da Silva and Tabarelli, 2000; Moran et al., 2009).

Combining systematic measures of 3D forest structure, such as those only recently available from spaceborne lidar (Dubayah et al., 2020), with analyses of functional traits is a promising avenue for understanding relationships between forest structure and adaptive morphologies of fauna for navigating vertical niches of tree canopies. In addition to advancing empirical analyses using ever expanding databases on species functional traits and ecosystem structure, future work could assess feedbacks between vertical niches, trophic interactions, dispersal, and ecosystem function using process-based ecosystem models that can incorporate vertical vegetation structure and functional interactions between organisms (Harfoot et al., 2014).

## To be Added Manually to Reference List

GBIF Secretariat (2021). GBIF Backbone Taxonomy. Checklist dataset https://doi.org/10.15468/39omei accessed via GBIF.org on 2021-11-12.

IUCN 2019. *The IUCN Red List of Threatened Species*. *Version 2019-1.18.* http://www.iucnredlist.org. Downloaded on 02 December 2019.

BirdLife International and Handbook of the Birds of the World (2018) Bird species distribution maps of the world. Version 2018.1. Available at http://datazone.birdlife.org/species/requestdis.

